## Supplementary material for "Polygenic architecture of flowering time and its relationship with local environments in the grass *Brachypodium distachyon*": FiguresS

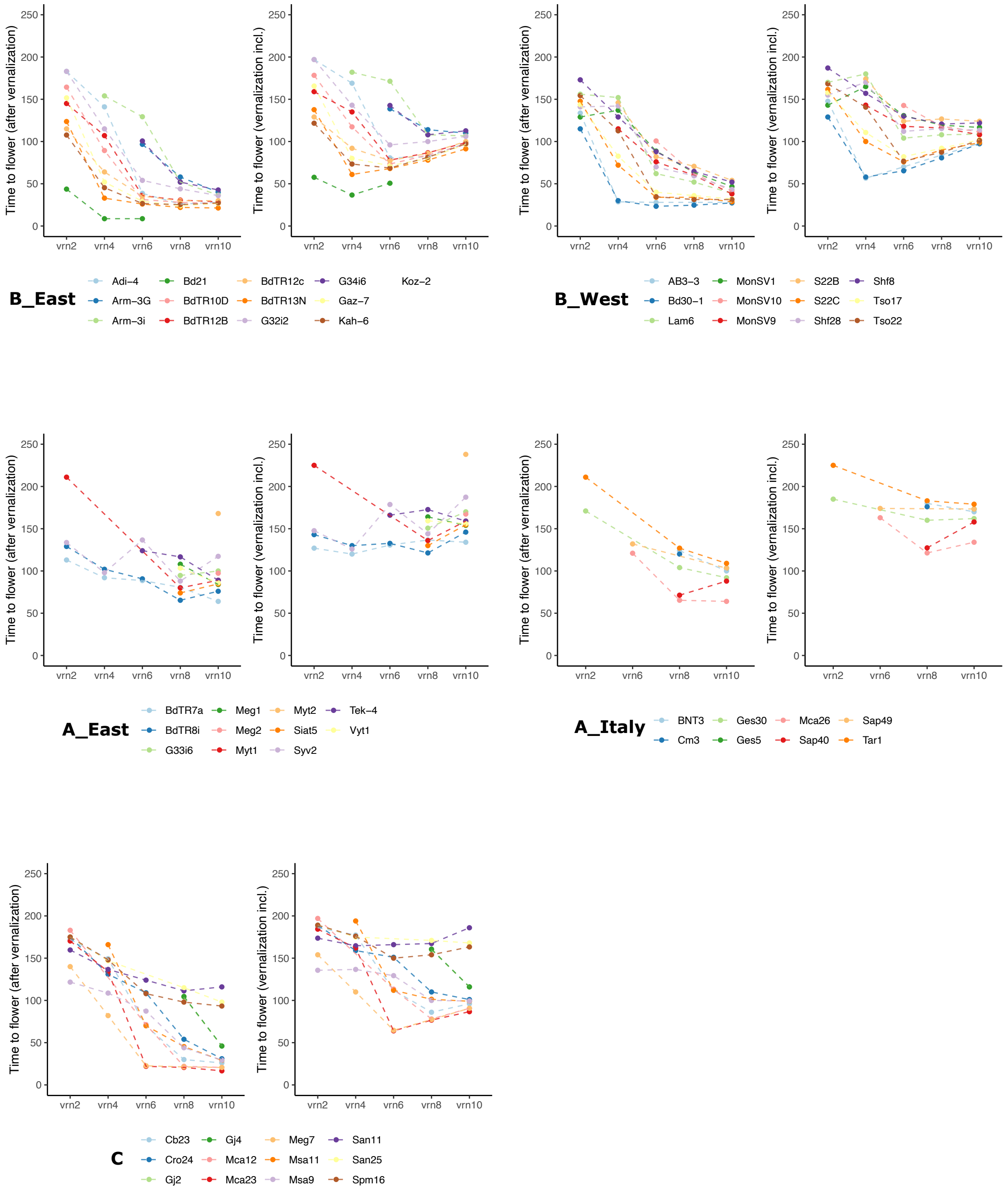


**Fig. S1:** Flowering time results**.** For each genetic clade, plots display the number of days given accessions (mean over replicates) take to flower after or including vernalization time.

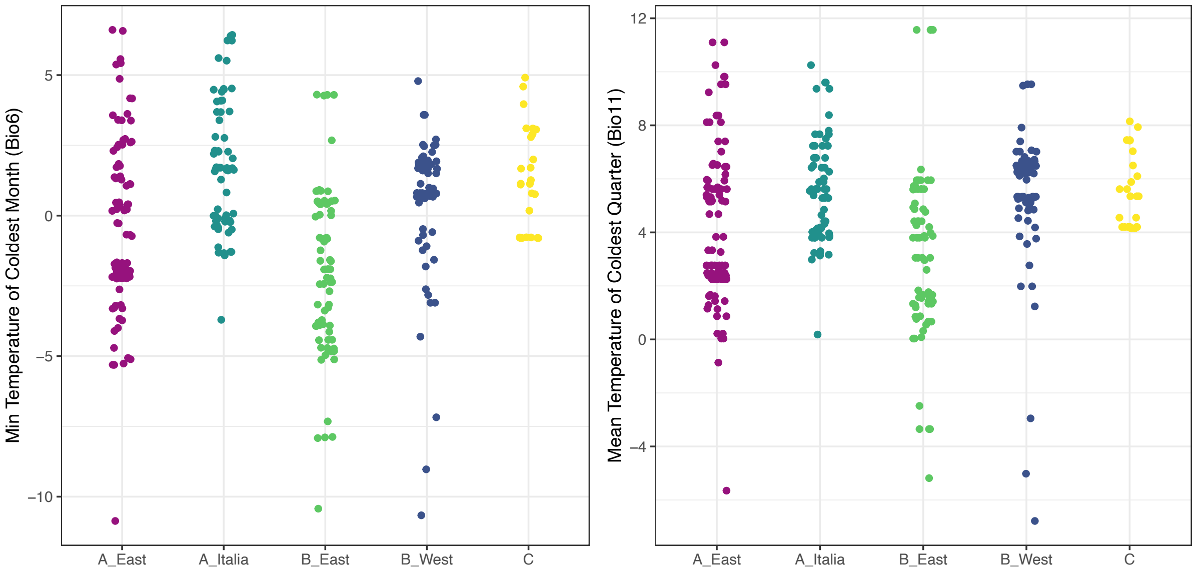


**Fig. S2:** Bio6 and Bio11 distribution across the 332 accessions of the diversity panel.

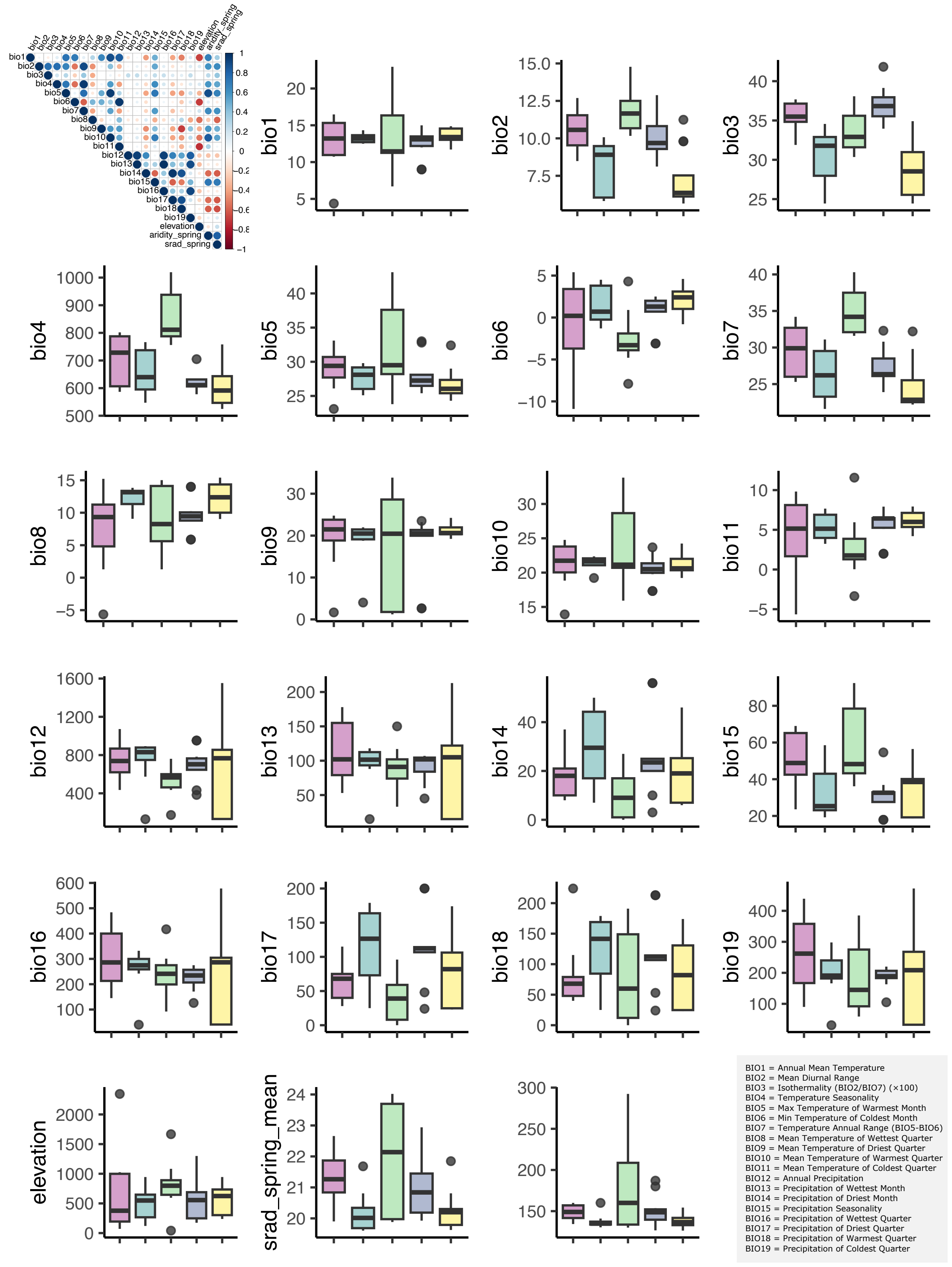


**Fig. S3:** Bioclimatic variable variation among the five genetic clades and 56 accessions chosen for the greenhouse experiment. The correlogram display le levels of correlation among variables.

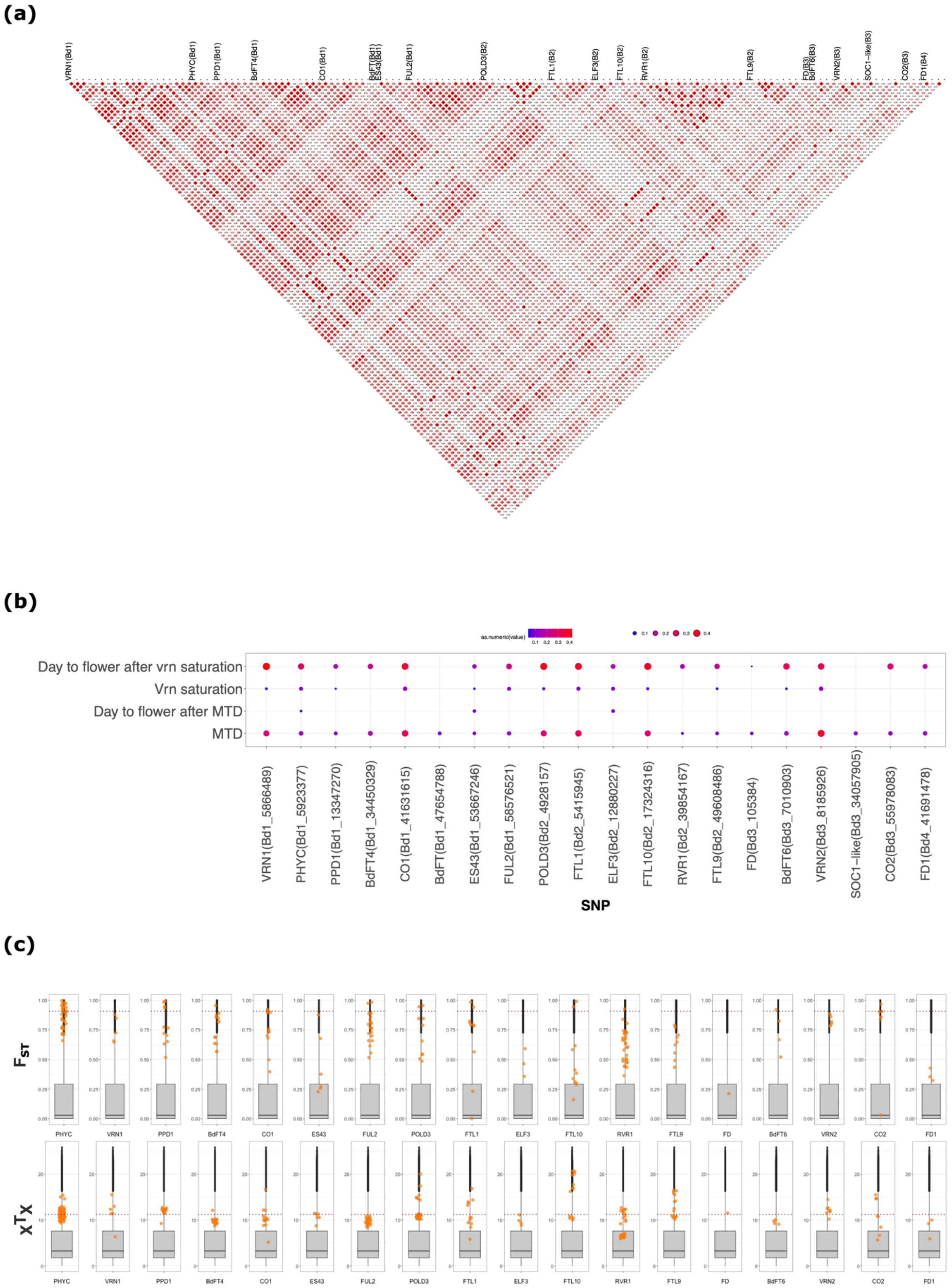


**Fig. S4:** Flowering time genes**.** (a) Linkage disequilibrium across the 20 flowering time genes associated with one of the flowering time-related trait (143 SNPs). The annotated SNP indicate the first SNP of each gene (b) RDA output for the 20 top SNPs associated flowering time-related traits. (c) $F_{ST}$ and $X^TX$ statistics computed in the entire diversity panel.

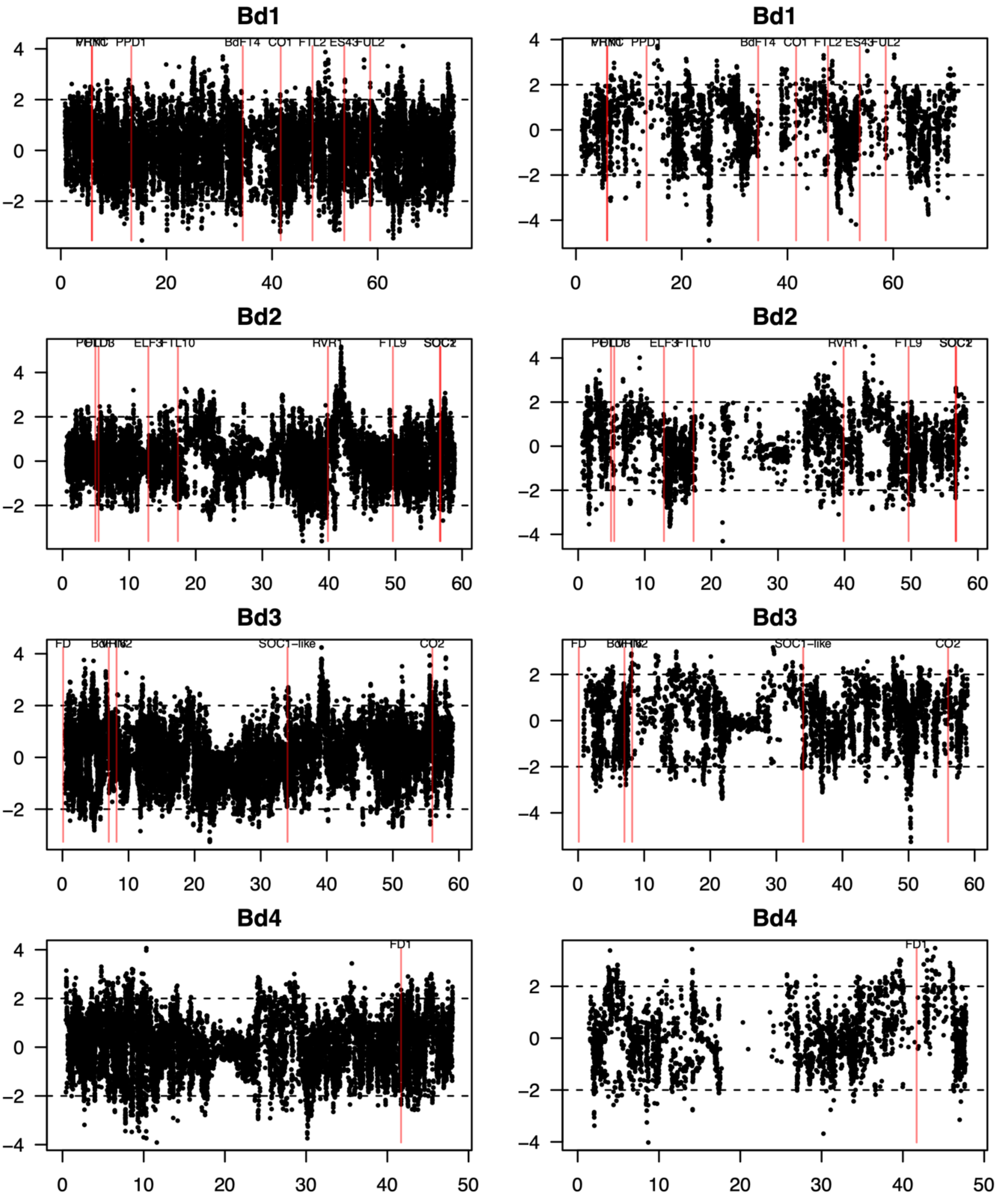


**Fig. S5:** integrated Haplotype Score (his) computed in the A (left panels) and B (right panely) lineages respectively.

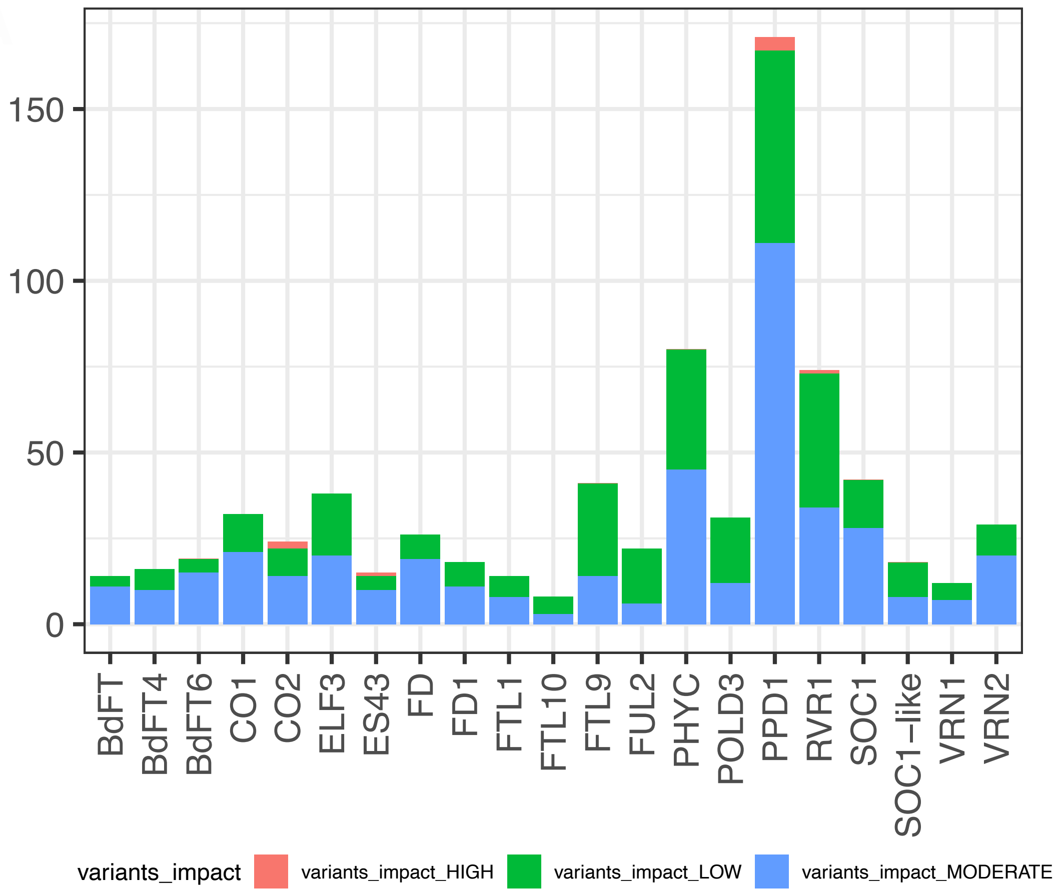


**Fig. S6:** SnpEff output for the flowering-time genes

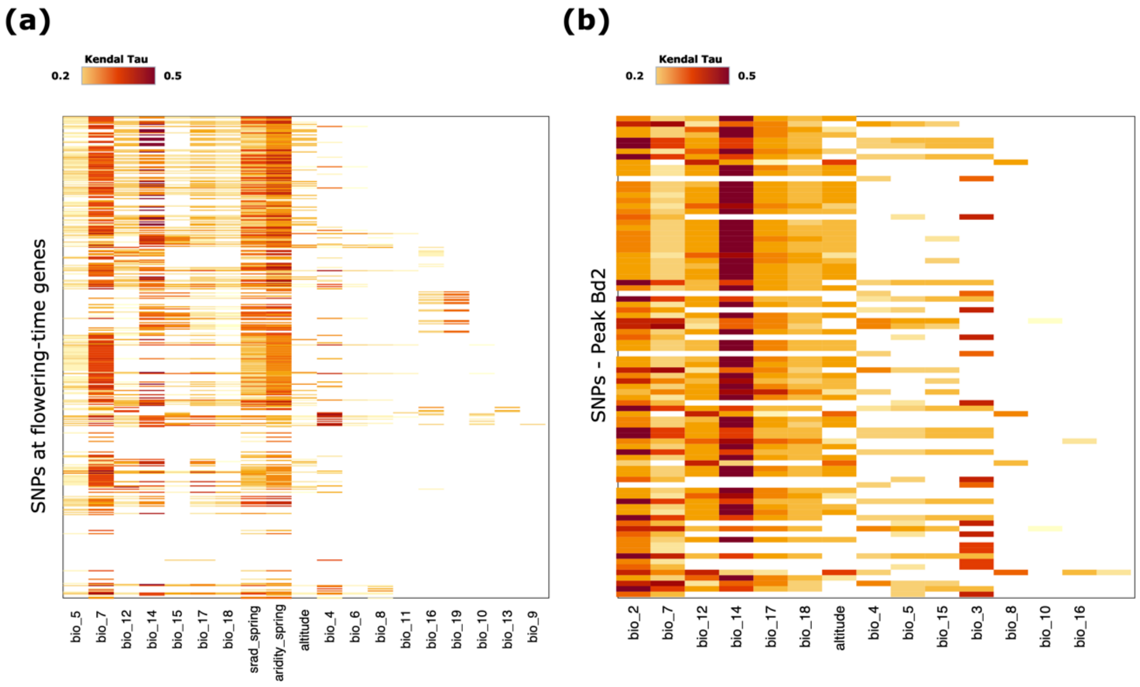


**Fig. S7:** Kendall correlation between SNP at AFT-genes and bioclimatic variables a) Heatmap displaying the association between SNPs at SFT-genes and bioclimatic variables. Variables not significantly associated with SNPs are not displayed b) Most significantly associated SNPs for the 9 SFT-genes showing a significant association with at least one bioclimatic variable c) Heatmap displaying the association between SNPs at GWAs peaks (outdoor experiment). Variables not significantly associated with SNPs are not displayed

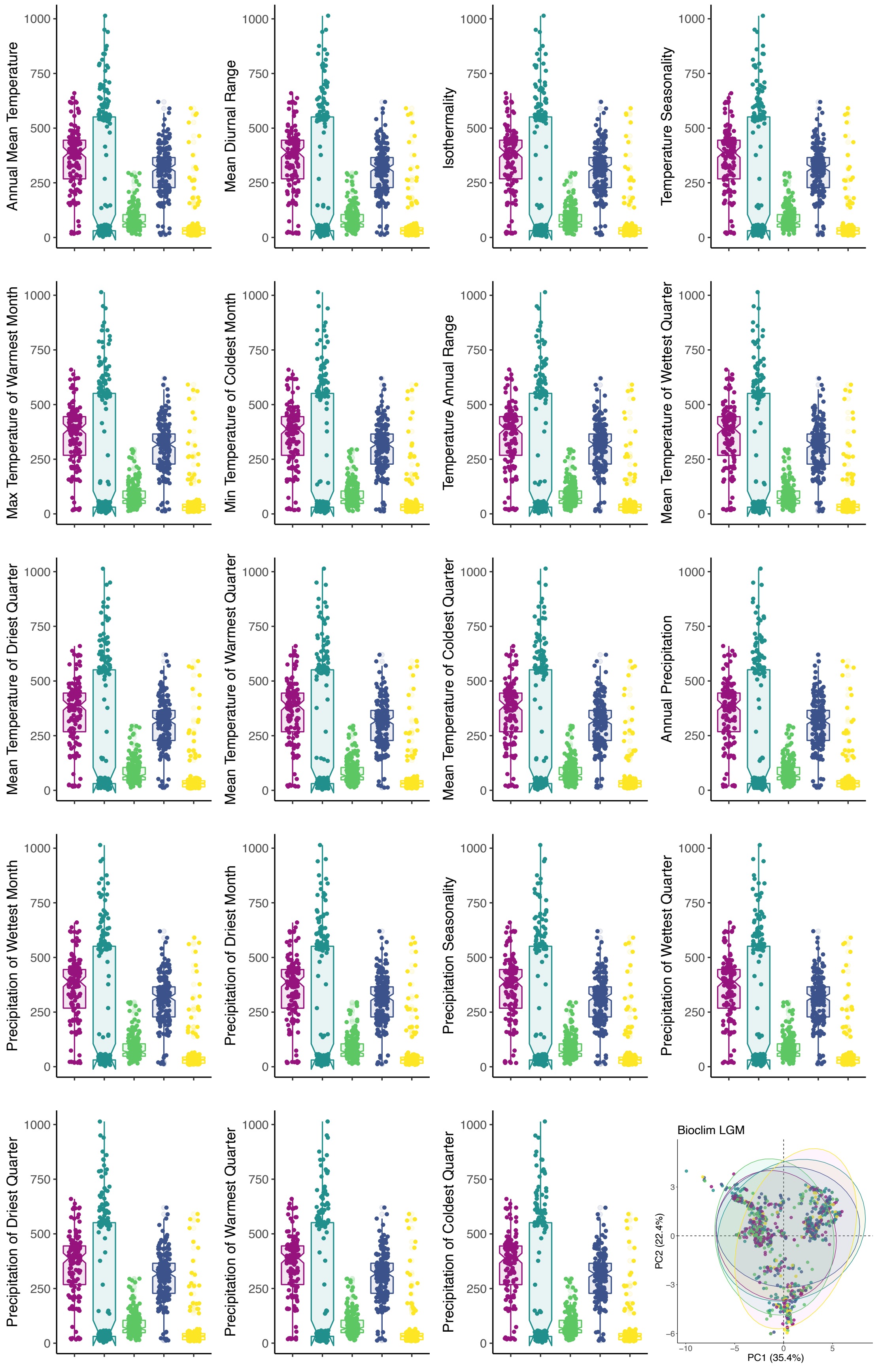


**Fig. S8:** Bioclimatic variables distribution over the five genetic clades during LGM. The PCA was performed using the 19 bioclimatic variables displayed by the boxplots.

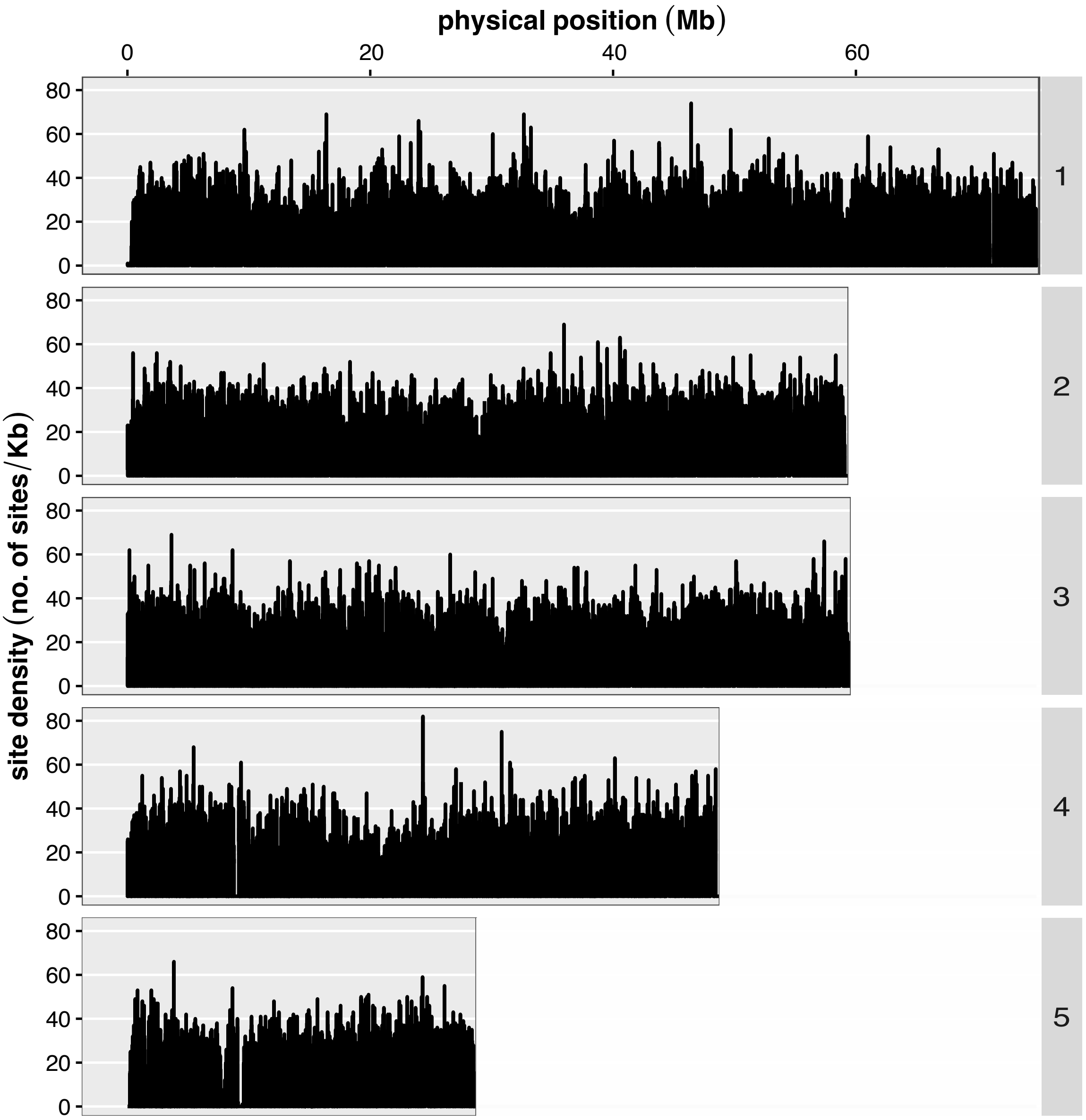


**Fig. S9: Marker density along the five chromosomes.** The plot displays the 2,266,225 SNPs kept for the GWAs
